# Hemodynamic Characteristics of Acute Hypersensitivity Reaction Induced by PEGylated Nanocomposite in Rats: Mechanism at the Nano-Bio Interface

**DOI:** 10.1101/2025.03.12.642931

**Authors:** Sin-Ting Ngo, Yun Cheng, Si-Yi Chen, Ming-Tsung Sun, Jau-Song Yu, Kun-Yi Chien, Hsi-Mei Chen, Yunn-Hwa Ma

## Abstract

**Background:** Polyethylene glycol (PEG)-modified nanocomposites may induce acute hypersensitivity reactions (HSR), including complement activation and hypotension, followed by tachyphylaxis with an unknown mechanism. We established a rodent model of acute HSR, and hypothesized that the formation of protein corona with a composition specific to PEGylated nanoparticles induces an acute and transient microvascular occlusion that entrains the hemodynamic effects.

**Methods:** Hemodynamic parameters of renal and cremaster vasculature were measured in anesthetized rats using ultrasonic flowmetry and laser speckle contrast imaging, respectively. Proteomic analysis on the hard corona of dextran-coated magnetic nanoparticles (MNP) with and without PEGylation was conducted after incubation of the nanoparticles with plasma from rats.

**Results:** PEG-MNP *iv*. induced a temporary reduction by approximately 30 mmHg in arterial pressure, with significant reduction in renal/cremaster blood flow and cardiac output, followed by tachyphylaxis and thrombocytopenia. PEG-MNP, but not pristine MNP, significantly increased renal vascular resistance with a reduction in the calculated cross-sectional area of renal vessels, suggesting microvascular occlusion. In contrast, the vasodilator acetylcholine decreased both blood pressure and vascular resistance before and after administration of PEG-MNP, suggesting an intact endothelium. Complement depletion by cobra venom factor induced a transient reduction in blood flow and prevented PEG-MNP-induced hemodynamic effects, suggesting an important role of complement activation. Proteomic analysis identified much higher complement proteins in the hard corona of PEG-MNP *vs*. MNP in plasma from rats; preexposure of rats to PEG-MNP or MNP *in vivo* greatly reduced plasma proteins with high affinity for PEG-MNP. The results suggest that complement depletion may mediate tachyphylaxis in response to the 2^nd^ dose of PEG-MNP.

**Conclusion:** PEGylated nanocomposites-induced complement activation in the protein corona may trigger the hemodynamic effects and subsequent pathophysiological responses in HSR of rats.

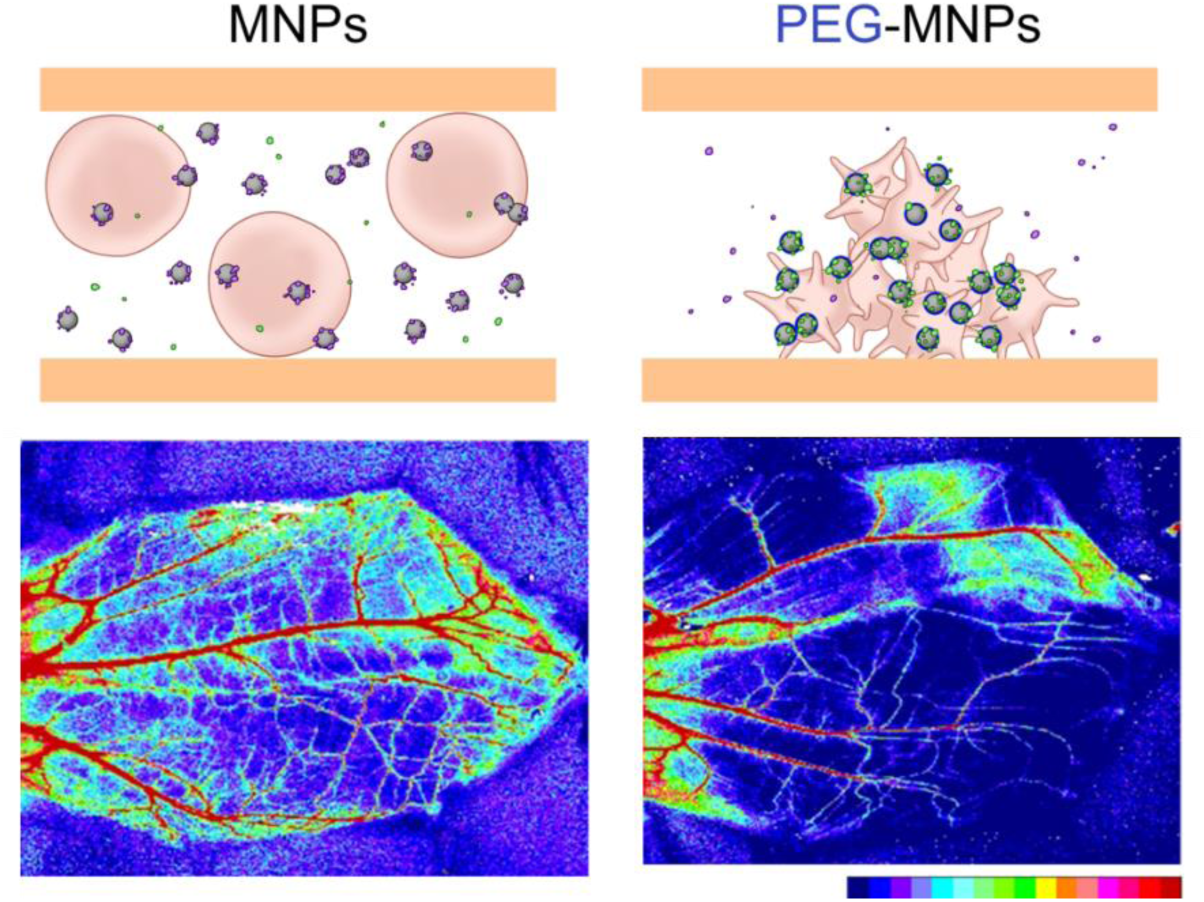

PEGylated magnetic nanoparticles (PEG-MNPs)-induced complement activation and platelet aggregation mediate the systemic increase in resistance of microvessels in rats.

---

Polyethylene glycol (PEG) is widely used as a surface modifier to increase the circulating half-life, stability, and therapeutic index of nano-biopharmaceuticals.^1–3^ Conjugation of PEG to therapeutic proteins or nanocarriers results in a larger hydrodynamic size, improves solubility, and provides steric hindrance to plasma proteins or blood cells that can cause clearance from the circulation.^1, 2^ Due to its good biocompatibility, PEG has been commonly used in the food and pharmaceutical industry, including most clinically approved nanomedicines.^4^ However, administration of PEGylated pharmaceuticals,^5–8^ including lipid-based mRNA COVID-19 vaccines,^9^ may induce acute hypersensitivity reactions (HSR). Such a reaction can lead to unpredictable lethal outcomes, especially in those with a history of severe allergy. Although premedication and desensitization may mitigate symptoms in most cases, HSR remains difficult to predict or prevent.^8, 10^

Nanobiopharmaceutical-induced HSR have been studied primarily in pigs for the past three decades,^8^ as pigs appear to be a sensitive model.^11^ Szebeni et al. have shown that acute HSR are associated with transient hypotension, increased pulmonary arterial pressure and resistance, decreased cardiac output (CO), thrombocytopenia, and up to 100-fold increase in plasma thromboxane B_2_, a stable metabolite of thromboxane A_2_ (TxA_2_).^12^ It was suggested that TxA_2_, a potent vasoconstrictor, may mediate cardiopulmonary distress;^13^ however, it is intriguing that the production of a vasoconstrictor should be associated with systemic hypotension. Although the observed cardiopulmonary hemodynamics appear to be similar to those of humans with HSR, tachyphylaxis occurred after the first injection in both pigs and rats with unknown mechanisms.

In addition to hemodynamic responses, HSR induced by nanobiopharmaceuticals are manifestations of complement activation-related pseudoallergy (CARPA).^4–7^ Activation of complement C3 and C5 entrains the secretion of secondary mediators, such as TxA_2_, by downstream effector cells.^6, 14^ Proteins in the complement system can be the main components in the protein corona on the surface of nanocomposites in circulation, and participate in modulation of the surface characteristics of nanocomposites, and thus their behavior/fate in circulation.^15–19^ Limited evidence suggests that complement activation may account for, at least in part, the hemodynamic effects associated with HSR in pigs^13^ and rats.^20^ It has been shown that the extent of complement activation is linearly associated with the amplitude of hypotensive effects in rats.^20^ However, how complement activation induces hypotension or tachyphylaxis remains to be elucidated.

For translation, nanosafety has become a pressing issue in the advancement of biomedical applications of nanomaterials;^21^ determination of the mechanisms underlying PEGylated nanomaterial-induced adverse effects appears to be critical in the design/development of nanomedicines. Recent studies indicate that a variety of nanoparticles can interact with vascular endothelial cells, induce an inflammatory response, and alter endothelial function.^22^ Whether PEGylated nanoparticles may exert an effect that contributes to hemodynamics is uncertain. Since superparamagnetic iron oxide nanoparticles, or MNPs, have been shown to have desirable biocompatibility and clinical potential,^23, 24^ we tested the hypothesis that PEGylated MNPs (PEG-MNPs) may induce acute HSR in anesthetized rats, which serves as a model for CARPA-associated acute hemodynamic responses. Since magnetic capture of PEG-MNPs may exhibit a size-dependent hemodynamic effect on microcirculation,^25^ a rat model was established with ultrasonic measurement of blood flow, allowing for a detailed analysis of hemodynamics in response to parenteral administration of PEG-MNPs. The results indicate that PEGylated MNPs induced a transient hypotensive effect with a decrease in tissue blood flow and an increase in peripheral vascular resistance (PVR), which were likely due to a transient microvascular occlusion associated with CARPA. The mechanisms of the hemodynamic responses may promote development of benchmarks in testing potential HSR of future nanomaterials.

## MATERIALS AND METHODS

### Materials

Dextran-coated magnetic nanoparticles (MNPs; nanomag^®^, 50 and 250 nm) with or without polyethylene glycol (PEG-600 or PEG-2000) were purchased from Micromod Partikeltechnologie (Rostock, Germany). If not specified, PEG-MNP refer to MNP coated with PEG-600. Acetylcholine (ACh) was purchased from Sigma-Aldrich (Burlingame, CA, USA). Polyethylene glycol (PEG) 1000 Da was purchased from Nanocs Inc. (NY, USA). Cobra venom factor was from Quidel Corporation (Athens, OH, USA). Sequencing grade trypsin was purchased from Promega (Madison, WI, USA), and other chemicals and solvents for proteomics were from Sigma-Aldrich (Burlingame, CA, USA).

### Methods

#### Hemodynamic measurements

The rat protocol was approved by the Institutional Animal Care and Use Committee of Chang Gung University, which is certified by the American Association for Accreditation of Laboratory Animal Care. Male Sprague Dawley rats (SD; 358 ± 3, n=70; BioLASCO Taiwan Co.) were anesthetized with Inactin^®^ (100 mg/kg) by intraperitoneal injection (*ip*). Body temperature was measured with rectal thermometry and maintained at 37℃ using a homeothermic blanket control unit (Harvard Apparatus Ltd.). The trachea is then cannulated to ensure a patent airway; while the urinary bladder was cannulated to keep patent urine flow. Blood pressure was monitored through carotid artery cannulation and a pressure transducer; heart rate (HR) was derived from the pulsatile blood pressure tracings. A mixture of 0.9% NaCl and 6% HEAS-steril^®^ (2:1) was continuously infused during (180 μl/min·kg) and after surgery (30 μl/min) into the right jugular vein to maintain plasma volume. The flows of the renal artery and abdominal aorta were monitored with an ultrasonic flow transducer (1 and 1.5 RB; Transonic System). ACh (1, 3 and 5 μg/kg), PEG (200 mg/kg) or MNP with or without PEG (randomly allocated; 5 or 10 mg/kg; 50 or 250 nm) was administered *via* the jugular vein. In another group of rats, cardiac output was measured according to previously described.^26^ Briefly, anesthetized rats were subjected to a thoracotomy with the trachea connected to a mechanical ventilator (Model 683; Harvard Apparatus Inc.), followed by placement of an ultrasound flow probe (2.5 PSB; Transonic System Inc.) on the ascending aorta of rats with a fluid supplement at a rate of 150 μl/min *via* jugular vein. The cross-sectional area analysis is detailed in the Supplemental Materials.

#### Rotational Thromboelastometry

After exposure to PEG-MNP (*iv*), the blood withdrawn from the anesthetized rats was subjected to thromboelastometry (ROTEM, Tem Innovations GmbH, Germany) as described previously.^27^ Coagulation was initiated with CaCl_2_ reagent (NATEM) or the diluted (1:100 in PBS) activators of extrinsic (EETEM)/intrinsic (INTEM) pathways of coagulation.

#### Cremaster microcirculation model

Male SD rats weighted 352 ± 8 g (n=10) were anesthetized with Inactin^®^ (100 mg/kg, *ip*). Body temperature was monitored using rectal thermometry and maintained at 37℃ using a homeothermic blanket control unit (Harvard Apparatus Ltd.). Cannulation procedures for the trachea, bladder, carotid artery, and jugular vein were similar as stated above. The cremaster microcirculatory preparation of rats was prepared on the basis of a previously described method. The layer of cremaster muscle was spread radially on a homemade stage to observe cremaster perfusion. PEG-MNP or MNP (10 mg/kg; 250 nm) was administered through the jugular vein while endothelial-dependent vasodilation was induced by topical Ach superfusion (0.1 or 0.2 mg/ml). Cremaster muscle was continuously superfused with physiological salt solution at room temperature to maintain equilibrium of the liquid layer on the muscle piece for a steady and continuous recording of blood flow in a real-time manner using the laser speckle contrast imaging technique (Moor Instruments).

#### Preparation of microparticle-free plasma

Rats anesthetized with isoflurane (3%) were subjected to cardiac puncture for blood collection with sodium citrate as anticoagulant. The whole blood was pooled and centrifuged at 1,000 × g (Soryall® RT 6000B) at 20℃ for 15 min within one hr of blood collection; the supernatant was then obtained and centrifuged at 3,720 × g (Backman Coulter Aventi® J-20XP) at 4℃ for 10 min to eliminate the cellular particles, including platelets. Lastly, the supernatant was centrifuged at 100,000 × g at 4℃ for one hour using an ultracentrifuge (Backman Coulter Optima^TM^ XE-90). Lipid suspension from the meniscus was removed to obtain microparticle-free plasma (MFP). The proteins in pristine rat MFP were quantified using Pierce’s BCA protein assay kit (Thermo Scientific) according to the manufacturer’s protocol.

#### Surface charge characterization

The Zeta potential measurements of the nanoparticles were determined using a Zetasizer (NanoZS90, Malvern Instruments) at 25℃. MNP versus PEG-MNP (4.8 μl) were thoroughly suspended in ddH_2_O or rat MFP (300 μl) and subsequently incubated at room temperature for 0 or 10 min, followed by immediate dilution and analysis of ddH2O.

#### Isolation of hard corona protein

MNPs or PEG-MNPs (PEG 600; 48 μg) were mixed with rat MFP (300 μl), incubated at 37℃ for 15 min using a digital dry bath, and manually mixed by pipetting every 3 min. A brief centrifugation at 10600 ×g for 1 min and magnetic separation (MagnaRack^TM^, Invitrogen) for 4 min allowed collection of particles with the protein corona, which was washed sequentially with 1× PBS (in mM: KCl 2.7, Na_2_HPO_4_.H_2_O 10, and KH_2_PO_4_ 1.8) containing 0.1% CHAPS and increased salinity (0.1, 0.2, 0.5 and 1 M NaCl), pH 7.4, to remove “soft corona” proteins. Then, the particles enriched with ‘hard corona’ proteins were incubated with SDS sample buffer at 100℃ for 10 min and subjected to one-dimensional sodium dodecyl sulfate polyacrylamide gel electrophoresis (SDS-PAGE) on a 10% Tris-Glycine gel. A PageRuler-prestained protein ladder (Thermo Scientific) as the molecular weight standard (10-170 kDa) was also run on the gels as a molecular weight standard (10-170 kDa). Finally, protein bands were detected and visualized by silver staining and scanned on ImageScanner III (GE Healthcare, IL, USA).

#### Liquid chromatography-mass spectrometry

The peptide mixtures were analyzed by an UltiMate 3000 RSLCnano HPLC system coupled with an Orbitrap Elite mass spectrometer (Thermo Fisher, Waltham, MA, USA). Concisely, the peptide mixtures were reconstituted in HPLC buffer A (0.1% formic acid in water), and loaded onto a trap column (Zorbax 300SB-C18, 0.3 × 5 mm; Agilent Technologies, Taiwan) at a flow rate of 20 μL/min with HPLC buffer A, and separated on an analytical column (ACQUITY UPLC column M-Class Peptide BEH C18, 0.1 × 100 mm, Waters Corporation). The peptides were subsequently eluted by the following gradient of ascending HPLC buffer B (0.1% formic acid in 100% ACN): 0-5 min, 5% B; 5-35 min, 5-30% B; 35-45 min, 30-45% B; 45-49 min, 45-95% B; 49-55 min, 95% B; 55-60 min, 5% B at a flow rate of 250 nL/min. The effluents were directly infused into the ion source of a mass spectrometer. The mass spectrometer was operated in positive ion mode using Xcalibur 2.0 software (Thermo Fisher). A data-dependent protocol containing one MS scan in the Orbitrap at a resolution of 60000 and 15 MS/MS scans in the linear ion trap for the 15 most abundant precursor ions was used to acquire data. The m/z scan range for MS was set as 400-2000 Da, and the cyclosiloxane ion signal at m/z 445.120025 was used as an internal standard for mass lock. Each precursor ion was selected once for MS/MS analysis and then excluded for 180 s.

#### Database search and protein quantification

The MS/MS spectra were searched with the Mascot Engine (Version 2.1, Matrix Science, Boston, MA, USA) against the Swiss-Prot protein sequence database (released in January 2020, selected for Rattus Norvegicus, 36181 entries) of the European Bioinformatics Institute using Proteome Discoverer 1.4 software. The mass tolerance for precursor and fragment ions was set as 10 ppm and 0.7 Da, respectively. Trypsin was established as the digestion enzyme and up to two missed cleavages were allowed. The searches were performed with the following parameters: Methylthiol (C) as a fixed modification; Acetyl (Protein N-term), Gln->pyro-Glu (N-term Q), Oxidation (M), Dimethyl (K), Dimethyl (N-term), Dimethyl: ^2^H_4_^13^C_2_ (K) and Dimethyl: ^2^H_4_^13^C_2_ (N-term) as variable modifications. The false positive identification rates were set to 1% as estimated by the Percolator algorism implanted in the Proteome Discoverer 1.4 software. The hard corona protein with at least 2 unique peptides was retained and the dimethyl heavy versus light ratio of each protein was log2 transformed followed by the removal of keratin prior to normalization. The normalization was then achieved by subtracting the median of all protein ratios. For proteins with a log2 ratio of PEG-MNP to MNP greater than 1 after normalization, they were considered to have a binding preference for PEG-MNP, while proteins with a log2 ratio of PEG-MNP to MNP below -1 after normalization were considered to be prone to MNP binding. The Mw and isoelectric point (pI) of all proteins were extracted using Protparam, a tool available in the ExPASy SIB Bioinformatics Resource portal.

#### Statistics

Values are means ± SEM. The effects were examined by factorial analysis of variance (ANOVA), followed by Duncan’s post hoc test. The student’s t test or the paired student’s t test was used as appropriate. *P*<0.05 was considered statistical significance.

## RESULTS

### PEG-MNP-induced renal hemodynamics and tachyphylaxis

Figure 1A illustrates representative changes in arterial pressure (AP), RBF, ABF, and vascular resistance (RVR/AVR) over time in response to consecutive doses of 250 nm PEG-MNP at 10 mg/kg. In this rat, the first dose of PEG-MNP reduced AP by 31 mmHg with a Tτ of 18 min, associated with a reduction in RBF and ABF by 6.2 and 10.2 ml/min, respectively. The results suggest that PEG-MNP-induced transient hypotension is associated with elevated systemic resistance. However, no significant changes were observed in all parameters in response to the second dose of PEG-MNP, suggesting a tachyphylaxis response.

**Figure 1.**
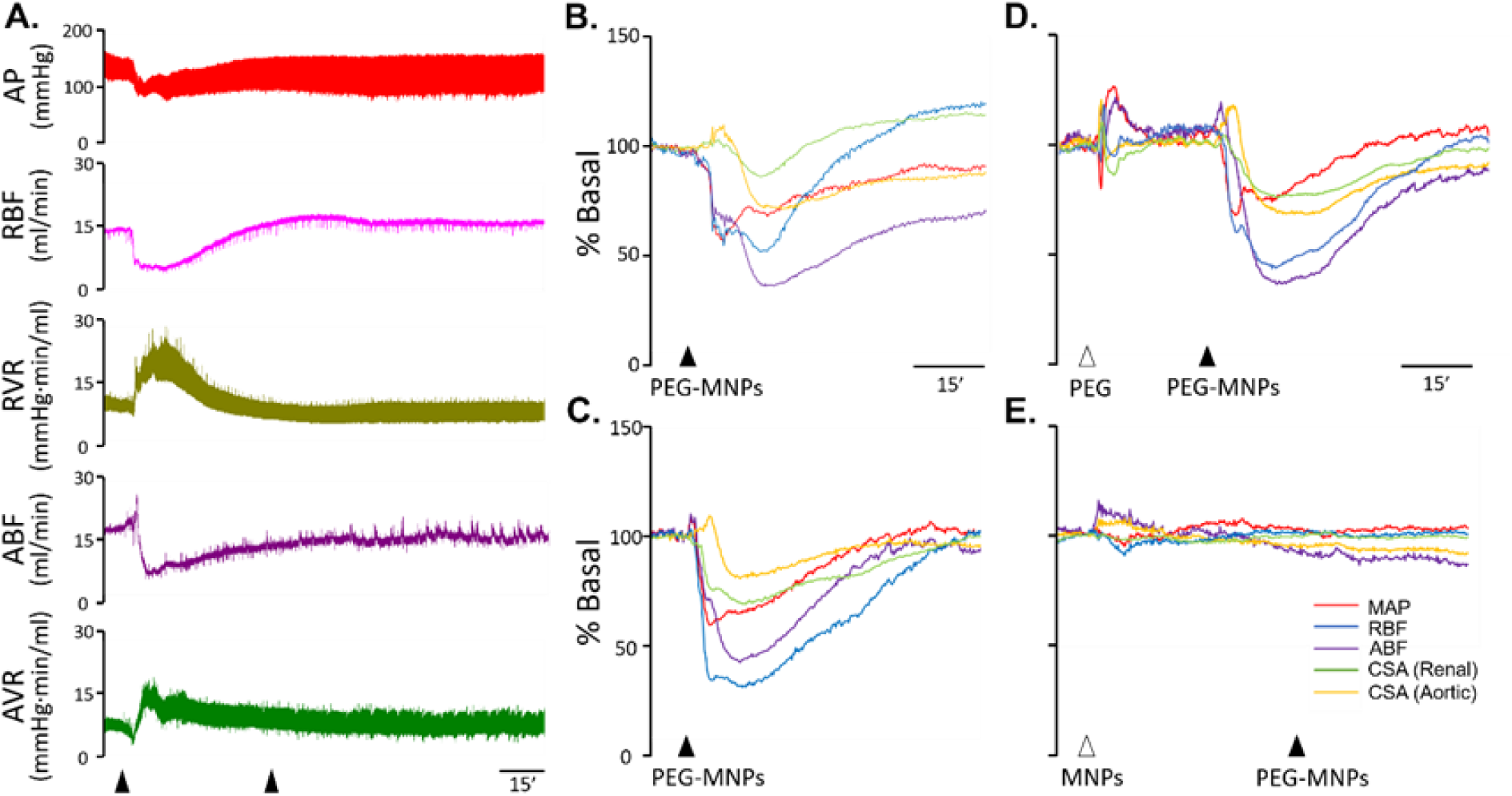
Representative hemodynamic responses to consecutive administration of PEG-MNP. (A) Representative hemodynamic tracings upon administration of consecutive doses of PEG-MNP (10 mg/kg), as indicated by the arrowheads. Cross-sectional area was calculated based on renal and aorta hemodynamic responses to 250 nm (B, D & E) and 50 nm (C) of PEG-MNP (10 mg/kg) *vs.* pre-treatment of free PEG (200 mg/kg; D) or MNP (250 nm, 10 mg/kg; E) as indicated by the arrowheads. Arterial pressure, AP; mean arterial pressure, MAP; renal blood flow, RBF; aortic blood flow, ABF; renal *vs.* aortic vascular resistance, RVR *vs.* AVR; renal *vs.* aortic cross-sectional area, RCSA *vs.* ACSA.

Figures 1B & C show an almost simultaneous reduction in mean arterial pressure (MAP), RBF, and ABF prior to the reduction of the calculated cross-sectional area (CSA) of the vasculature in response to PEG-MNP of 250 and 50 nm, respectively. It is noticed that there is often a transient increase in ACSA before a profound reduction response occurred; whereas RCSA did not exert such a transient increase. In general, similar patterns of responses are observed upon administration of 250 *vs.* 50 nm PEG-MNPs.

To determine whether PEG or the iron oxide particles may be responsible for the hemodynamics/tachyphylaxis of PEG-MNPs, PEG or MNPs were administered prior to PEG-MNPs. Figure 1D shows that PEG-MNP induced a reduction in MAP by 33 mmHg after pretreatment with PEG, which was associated with a decrease in RBF and ABF by 6.7 and 12.4 ml/min, respectively, in addition to a decrease in RCSA and ACSA. After pretreatment with PEG, the hypotensive effect induced by PEG-MNP became relatively transient with a Tτ of 380 s. On the contrary, hypotensive effects and associated changes in all hemodynamic parameters induced by PEG-MNP were completely abrogated by pretreatment with MNP (Figure 1E). In these rats, acetylcholine (Ach)-induced vasodilation was not significantly different before and after MNP or PEG-MNP administration, suggesting intact endothelial function after administration (Suppl. Figure 1). Suppl. Table 1 shows that without pretreatment, PEG-MNP reduced blood pressure by 31 mmHg, which was associated with a decrease in RBF and ABF and an increase in RVR and AVR, as illustrated above. However, such effects were not different from those parameters after PEG pretreatment. Because MNP pretreatment also abrogated PEG-MNP-induced hemodynamics, the common surface characteristics of MNPs and PEG-MNPs may be responsible for tachyphylaxis in rats.

Figure 2A illustrates that 250 nm PEG-MNP induced an average hypotensive effect of 31 mmHg, which was associated with a significantly reduced RBF and ABF by 57% and 32%, respectively, but increased RVR and AVR. However, the hemodynamic effects on MAP, RBF, and ABF were greatly reduced in response to the second dose of PEG-MNP, suggesting that tachyphylaxis occurred in all hemodynamic parameters. On the contrary, MNPs did not exert significant effects on MAP, RBF, or ABF. Furthermore, the first dose of PEG-MNPs, but not MNPs, induced a significant increase in RVR and AVR, followed by tachyphylaxis. A similar pattern of hemodynamic responses to 50 nm PEG-MNP *vs.* MNP at the same dose was also observed (Figure 2B). The basal hemodynamic parameters of the aforementioned groups remained relatively stable and did not show significant differences, as presented in Suppl. Table 2. A similar pattern of hemodynamic effects was observed in response to 5 mg/kg of PEG-MNP *vs.* MNP (250 nm); nevertheless, indomethacin (2 mg / kg) did not exhibit an effect on the hemodynamic effects induced by PEG-MNP (Suppl. Figure 2). Therefore, our results do not support a role of TxA_2_ in the effects of PEG-MNP. Furthermore, PEG-MNP (250 nm, 10 mg/kg; *iv*)-induced a significant reduction in MAP by 25%, which was associated with a 47% and 49% decrease in cardiac output (CO) and stroke volume (SV) from a baseline of 64 ± 5 ml/min and 171 ± 11 μl/beat, respectively (n=5); whereas no difference in HR was observed.

**Figure 2.**
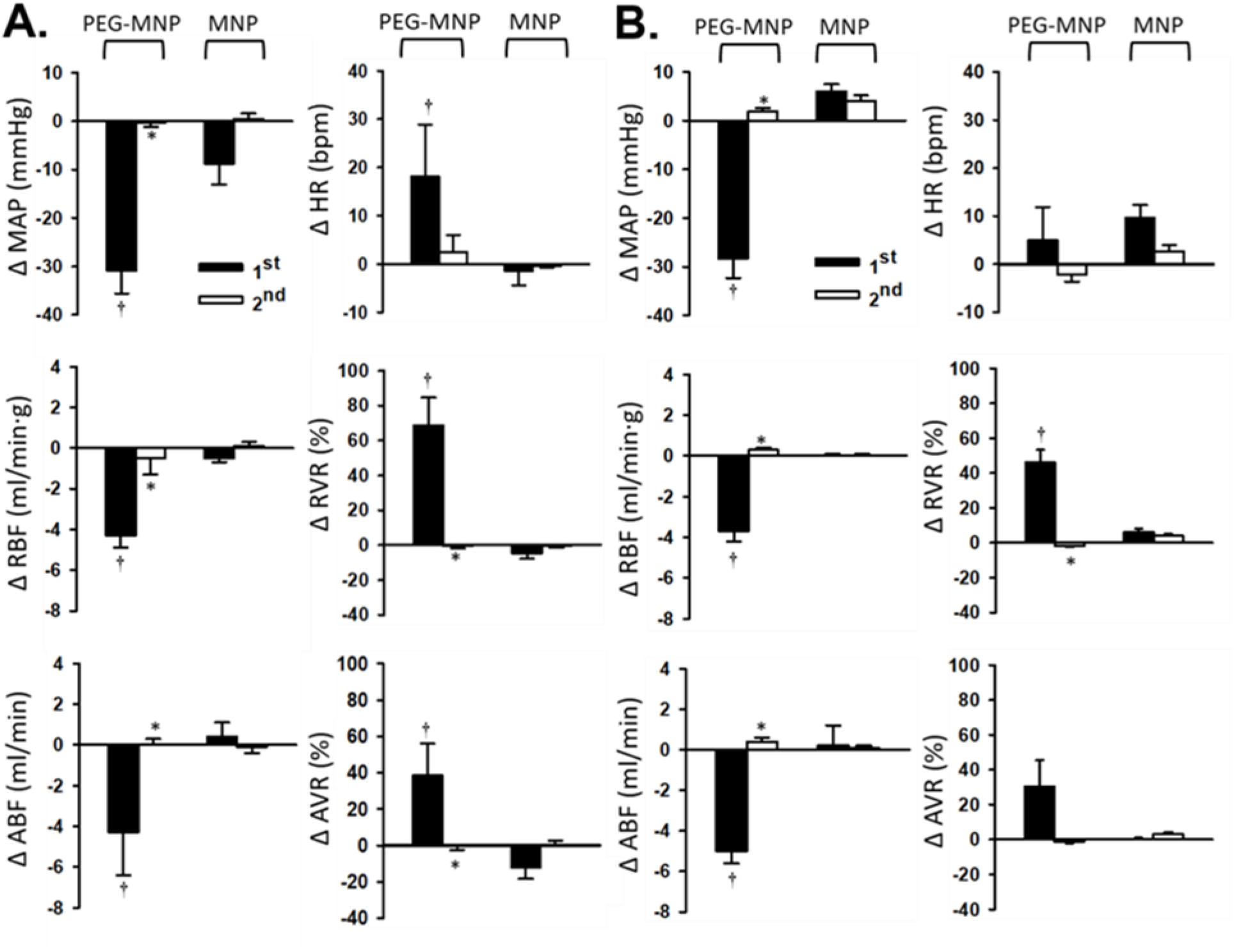
Analysis of tachyphylaxis associated with transient hypotension induced by PEG-MNP in anesthetized rats. Each rat was iv administered with two consecutive doses (10 mg/kg) of MNP or PEG-MNP with a size of 250 nm (A) or 50 nm (B), respectively (n=7-10). *†, P<0.05 compared with the 1^st^ dose and MNP, respectively. The results are representative of Protocols 3, 5, 6, and 7 in Suppl. Table 2. Mean arterial pressure, MAP; renal blood flow, RBF; renal vascular resistance, RVR; heart rate, HR; aortic blood flow, ABF; aortic vascular resistance, AVR.

### Hematology and histology analysis

Following *iv* PEG-MNP, blood was obtained through cardiac puncture of the rat for hematology analysis. Table 1 shows a significant decrease in platelet count by 40% and 50% in response to MNPs conjugated with PEG600 *vs.* PEG2000, respectively. In addition, the WBC count was reduced by PEG2000-MNPs, but not PEG600-MNPs. Thromboelastometry analysis (Suppl. Figure 3) on the coagulation activity of whole blood from rats exposed to PEG-MNPs indicates no difference in clotting time was observed in all groups. However, a decreased alpha angle was observed with PEG600-MNPs, and a decreased amplitude and maximum clot firmness were observed with both groups of PEG600-MNPs and PEG2000-MNPs. These effects on clot formation are consistent with thrombocytopenia in response to PEGylated nanocomposites in Table 1. At the end of some experiments, the kidney, lung, and liver were perfused and collected for histology analysis with Prussian blue staining. Suppl. Figure 4 illustrates that there is no retention of PEG-MNPs in the lungs or kidneys, as in the saline group, at the end of the experiments. In contrast, PEG-MNPs were found in the liver, as the liver is usually the major excretion organ. Nevertheless, no significant morphology change was observed in any organs studied after exposure to PEG-MNPs at the time studied.

**Table 1.** Hematology analysis after rats receiving MNPs with immobilized PEG600 *vs.* PEG2000.

|  | Saline | PEG600-MNP | PEG2000-MNP |
| --- | --- | --- | --- |
| n | 6 | 6 | 8 |
| Platelets (10<sup>3</sup>/uL) | 1057 ± 137 | 631 ± 93* | 534 ± 88* |
| RDW (%) | 13 ± 0.4 | 13 ± 0.4 | 13 ± 0.3 |
| MCHC (k/uL) | 32 ± 0.1 | 31 ± 0.4 | 31 ± 0.3 |
| MCH (pg) | 20 ± 0.3 | 20 ± 0.3 | 20 ± 0.3 |
| MCV (fL) | 64 ± 1.1 | 64 ± 1.0 | 64 ± 1.0 |
| Hematocrit (%) | 47 ± 1.4 | 47 ± 2.4 | 48 ± 1.5 |
| Hemoglobin (g/dL) | 15 ± 0.5 | 14 ± 0.7 | 15 ± 0.5 |
| RBC (10<sup>6</sup>/uL) | 7 ± 0.3 | 7 ± 0.4 | 8 ± 0.3 |
| WBC (10<sup>3</sup>/uL) | 7 ± 0.8 | 9 ± 1.0 | 6 ± 0.7<sup>#</sup> |

Two consecutive doses of PEG600-MNP, PEG2000-MNP (250 nm, 5 mg/kg) or saline (<200 µL) were administered intravenously to the rats prior to cardiac puncture to obtain blood for analysis. RDW, red cell distribution width; MCHC, mean corpuscular haemoglobin concentration; MCH, mean corpuscular haemoglobin; MCV, mean corpuscular volume; RBC, red blood cell; WBC, white blood cell. \*,<sup>#</sup>, $P < 0.05$ compared to saline and PEG600, respectively.

### PEG-MNP-induced cremaster hemodynamics and tachyphylaxis

A rat cremaster muscle model was used to visualize the effects of PEG-MNP on microvascular perfusion. Laser speckle contrast imaging demonstrates a significant decrease in cremaster tissue perfusion upon exposure to the first, but not the second, dose of PEG-MNP (10 mg/kg; Figure 3A). In contrast, the first or second dose of MNP did not induce such an effect (Figure 3B). Figure 3C illustrates that the first, but not second, dose of PEG-MNP induced a transient reduction in tissue perfusion in all vessels studied in the same preparation over time. The quantitative effects of the first and the second doses of PEG-MNP on the tissue perfusion (n=5) are illustrated in the insert of Figure 3C. In these rats, consecutive doses of Ach induced an increase in tissue perfusion before and after PEG-MNPs, suggesting that the endothelium of the cremaster microvasculature remained intact after administration of PEG-MNP.

**Figure 3.**
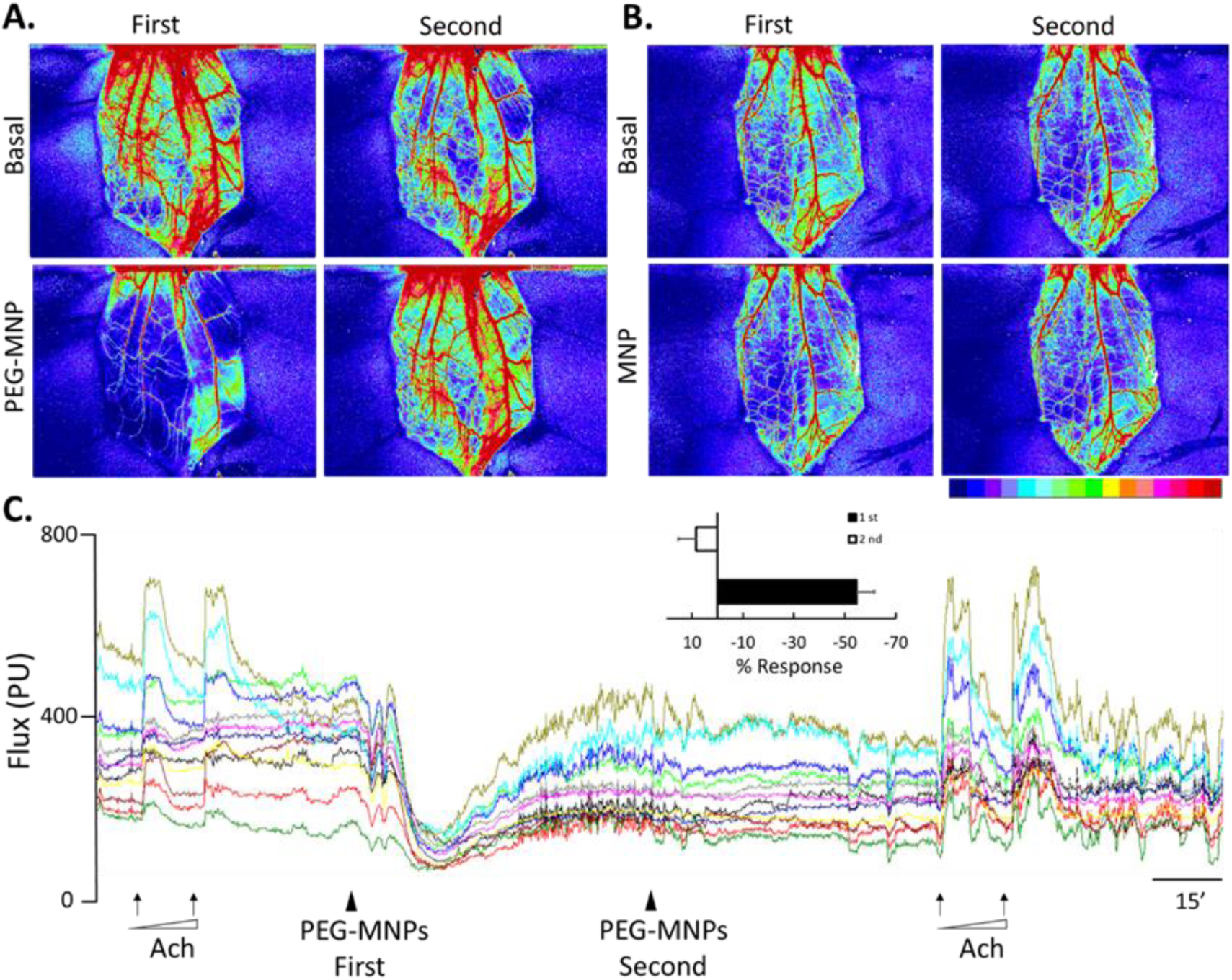
Tissue perfusion of cremaster microcirculation in response to consecutive doses of PEG-MNP vs. Ach. Laser speckle contrast images (A, B) and Flux with 12 ROIs (arbitrary unit PU) of 4 major vessels as a function of time (C) before and after two doses of PEG-MNPs *vs.* MNPs in the representative rats were recorded. Acetylcholine (Ach; 0.1 and 0.2 mg/ml) was superfused on the cremaster muscle as indicated by the arrows; PEG-MNPs (250 nm; 10 mg/kg) was administered iv as indicated by the arrowheads. The insert is quantitative results of the first and second doses of PEG-MNP in 5 rats studied.

### Complement depletion by the cobra venom factor

It is well known that cobra venom factor (CVF) activates and depletes the complement system in various experimental models.^28^ Figure 4A illustrates representative hemodynamic tracings in response to 5 IU/kg of CVF, with an increase in AVR/RVR and a decrease in ABF/RBF; pretreatment with CVF *iv* completely blocked changes in all hemodynamic parameters induced by PEG-MNP, but not Ach. Furthermore, 20 IU/kg of CVF induced greater changes in AP, RBF, and ABF, suggesting that CVF complement activation induces hemodynamic effects in a dose-dependent manner (Figure 4B).

**Figure 4.**
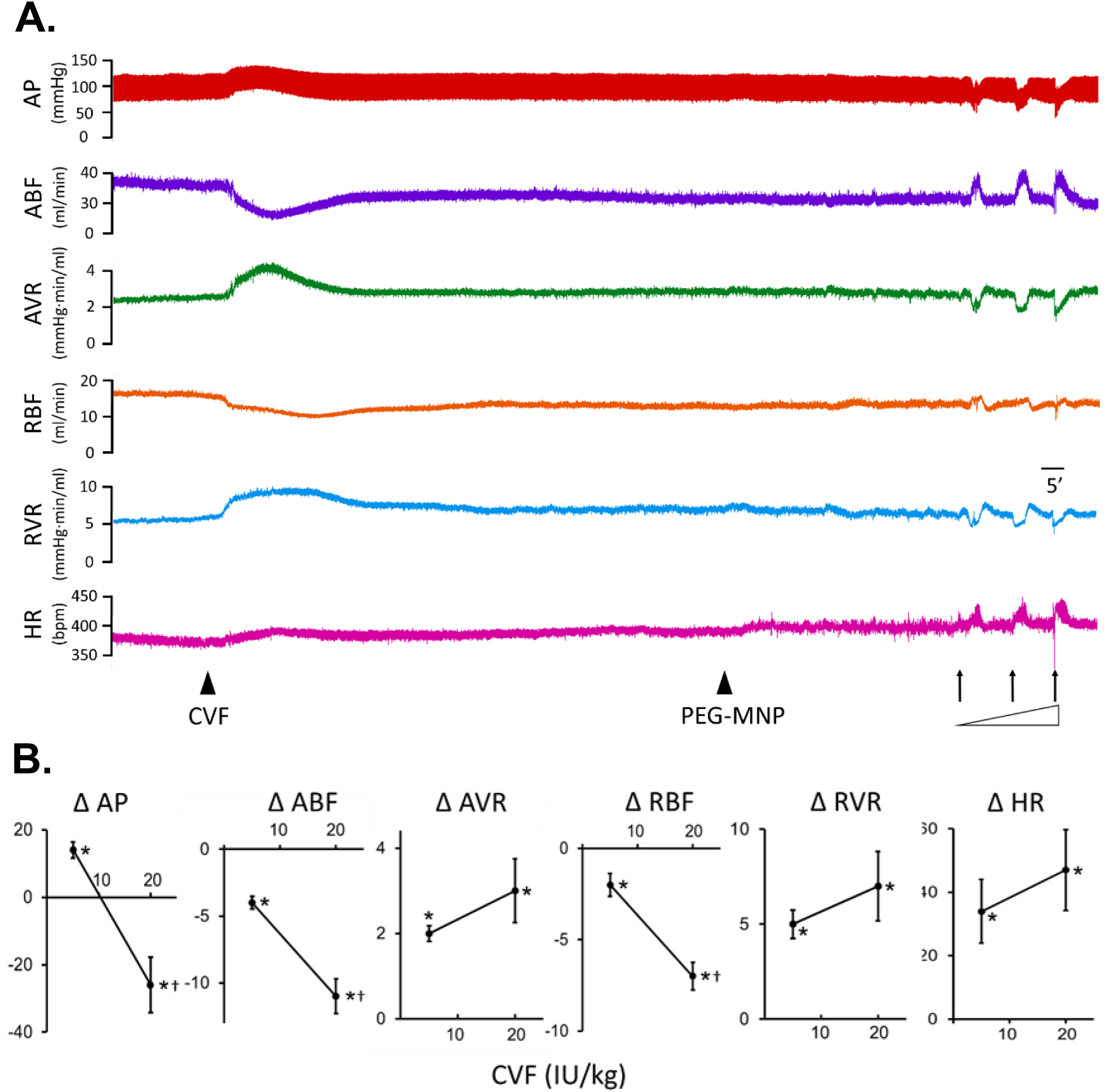
Pretreatment of cobra venom factor completely blocked PEG-MNP-induced hemodynamic effects in all rats studied. (A) Representative hemodynamic effects induced by *iv* administration of cobra venom factor (CVF; 5 IU/kg) followed by PEG-MNP (5 mg/kg) and acetylcholine (1, 3 and 5 mg/kg as indicated by arrows) were monitored in an anesthetized rat. (B) Does-dependent changes in hemodynamic responses to 5 *vs.* 20 IU/kg of CVF, with n=8 *vs.* 5, respectively. *, *P*<0.05, compared with the corresponding basal levels; †, *p* < 0.05, compared with the corresponding values of CVF 5 IU/kg. Arterial pressure, AP; aortic blood flow, ABF; aortic vascular resistance, AVR; renal blood flow, RBF; renal vascular resistance, RVR; heart rate, HR.

### Proteomic composition of the hard corona of PEG-MNs vs. MNP

To determine the mechanism underneath PEG-MNP-induced hemodynamic effects, nanoparticle-protein interaction was simulated *in vitro*. After incubation with rat plasma, the MNPs *vs.* PEG-MNPs were subjected to magnetic separation and the composition of the hard corona was determined, classified, and compared by proteomic analysis (Suppl. Figure 4). Table 2 shows that the predominant proteins in the hard corona of PEG-MNP *vs.* MNP were complement proteins, including complement C6, C3, C8, C9, and C4. Proteins with medium (M) or high (H) binding preference for both MNPs and PEG-MNPs were classified on the basis of their molecular weight (Mw) and isoelectric point (pI). Low Mw proteins with Mw < 30 kDa and Mw 30-50 kDa were equally enriched (37.5%) and together constituted the majority of solid binding proteins of MNP, while high Mw proteins with Mw > 100 kDa were found to be the most enriched proteins, constituted 50% of tightly bound proteins in the hard corona of PEG-MNP (Figure 5A). Furthermore, proteins with a strong binding preference for MNPs mainly exhibited pI > 9, in contrast to PEG-MNPs predominantly with a pI of 5-7 (Figure 5B).

**Figure 5.**
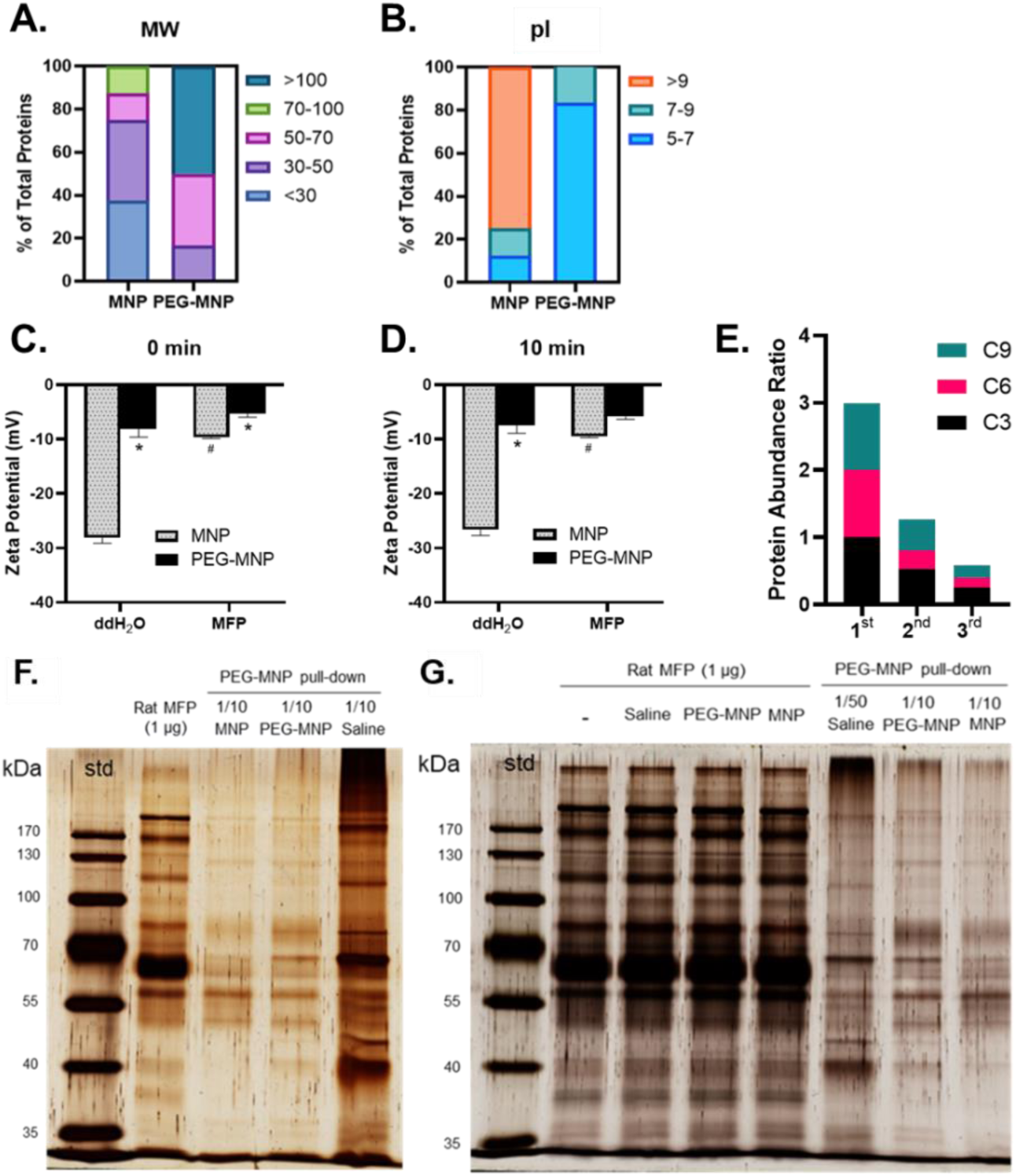
Proteomic profile of hard corona proteins on MNP *vs.* PEG-MNP. Distribution of proteins in the hard corona of MNP or PEG-MNP with medium or high abundance (Table 2) based on calculated molecular weight (MW; A) and isoelectric point (pI; B). (C) ζ-potential of MNPs *vs.* PEG-MNPs was measured in ddH_2_O or rat MFP after 10 min equilibration. (D) Hard corona proteins enriched by PEG-MNPs were derived from plasma of rats pre-treated with 10 mg/kg (*iv*) of MNPs, PEG-MNPs, or saline. The enriched plasma proteins were further separated with a 10% tris-glycine gel and visualized *via* silver staining (E & F). PEG-MNP pull-downs (1/10 of total enriched proteins) were compared to rat MFP (1 μg) without cannulation. Mean ± SEM, n=3; *, # *p* < 0.05, compared with the corresponding MNPs and ddH_2_O groups.

**Table 2.** Characteristics of hard corona proteins with high affinity for MNPs with or without PEGylation.

| Rank | Protein Name | PEG-MNP/<br>MNP<br>(Log 2 ratio) | Abundance | pI | KDa |
| --- | --- | --- | --- | --- | --- |
| PEG-MNP/MNP > 1 |  |  |  |  |  |
| 1 | Complement component C6 | 3.4 | H | 6.6 | 105.1 |
| 2 | Complement C3 | 3.3 | H | 6.1 | 186.5 |
| 3 | Complement component C8 beta chain | 2.5 | H | 8.4 | 66.7 |
| 4 | Complement component C9 | 2.0 | H | 5.5 | 62.3 |
| 5 | Pulmonary surfactant-associated protein I | 1.4 | M | 6.8 | 37.6 |
| 6 | Complement C4 | 1.0 | M | 7.0 | 192.2 |
| PEG-MNP/MNP < -1 |  |  |  |  |  |
| 1 | 40S ribosomal protein S9 | -3.6 | H | 10.7 | 22.6 |
| 2 | 60S ribosomal protein L7 | -2.8 | H | 10.9 | 30.3 |
| 3 | Platelet factor 4 | -2.6 | H | 9.6 | 11.3 |
| 4 | 60S ribosomal protein L7a | -2.5 | H | 10.6 | 30.0 |
| 5 | Kininogen-1 | -1.9 | M | 6.3 | 70.9 |
| 6 | 60S ribosomal protein L6 | -1.6 | M | 10.7 | 33.6 |
| 7 | Histidine-rich glycoprotein | -1.6 | M | 7.8 | 59.1 |
| 8 | 60S ribosomal protein L14 | -1.3 | M | 11.1 | 23.3 |

Proteins enriched by PEG-MNPs and MNPs at levels greater than a twofold ratio (PEG-MNP/MNP > 1 or < -1) show their predispositions to PEG-MNPs and MNPs binding, respectively. Enriched proteins by PEG-MNPs or MNPs at levels greater than a fourfold ratio (PEG-MNP/MNP > 2 or < -2; shaded areas) indicate their particular binding preference to PEG-MNPs and MNPs accordingly. H, proteins with high abundance; M, proteins with medium abundance.

Furthermore, the zeta potential of MNP was greatly reduced in rat plasma compared to that in DDW at 0 and 10 min, whereas the zeta potential of PEG-MNP in rat microparticle-free plasma (MFP) was not considerably different from that in DDW at 0 min and 10 min. Lastly, the difference in the averaged zeta potential between MNPs and PEG-MNPs in rat plasma was, although slightly, distinct at 0 min; however, no difference was observed between MNPs *vs.* PEG-MNPs in plasma at 10 min (Figure 5C & D).

To determine whether complement proteins in rat plasma can be depleted by repetitive exposure, PEG-MNPs were added to the supernatant plasma after magnetic separation for the second and third incubation, followed by proteomic analysis (Suppl. Figure 6). The results indicate that the abundance of complement C9, C6, and C3 protein in the PEG-MNP pulldown of the corona decreased with the second and third pulldown (Figure 5E).

### Interaction of rat plasma proteins from rats pre-exposed to PEG-MNPs

The hard corona proteins of PEG-MNPs were obtained from incubation of PEG-MNPs with plasma from rats with *iv* administration of vehicle, MNPs or PEG-MNPs and subjected to wash and electrophoresis (Suppl. Figure 7). A 10% protein fraction (1/10) from each sample of PEG-MNP extraction demonstrates that plasma derived from rats without exposure to nanoparticles (saline as a vehicle) exhibits the highest protein adsorption, while the amount of PEG-MNP extraction from rat plasma pre-exposed to MNP *vs.* PEG-MNP was comparable, but with the difference in patterns (Figure 5F). Although no difference in protein profiles was observed between rat MFP obtained after exposure to saline, PEG-MNP *vs.* MNP, PEG-MNP pulldown exhibited different protein profiles among the three treatments (Figure 5G), suggesting that plasma protein consumption with pre-exposure of PEG-MNP may occur *in vivo*.

## DISCUSSION

PEG-MNP-induced hemodynamic effects in rats are associated with complement activation, tachyphylaxis and thrombocytopenia, which are comparable to acute HSR induced by PEGylated nanomedicine in humans. With the rat model, we first report an increase in peripheral vascular resistance (PVR) associated with HSR-induced hypotension; therefore, PEGylated MNP-induced-hypotension is unlikely mediated by a direct vasodilator effect. Since heart rate was not altered, PEG-MNP-induced significant reduction in cardiac output (CO) was likely mediated by a reduced venous return due to transient occlusion of microvessels with platelet aggregates. The results reveal that a depletion of the complement proteins and platelets appears to cause tachyphylaxis induced by PEG-MNPs. The finding may shed the light on future prevention or treatment of the adverse effects induced by PEGylated nanomedicine.

CVF induced a transient reduction in RBF of rats in this and previous studies.^29^ In the current study, CVF completely blocked the hemodynamic effects of PEG-MNP in all rats studied; therefore, complements appear to play a crucial role in mediating the hemodynamic effects of PEG-MNP. In addition, complement inhibition may completely block complement opsinization of various nanoparticles.^30^ Although we could not rule out proteins other than complements may also participate in acute HSR, these findings suggest that modulation of the complement system may be an potentially effective strategy in control of acute HSR induced by nanomedicines.

Since complement components and activation products were found in the hard corona of PEG-MNPs *vs*. MNPs, complement activation may occur on the surface of PEG-MNPs that triggers a series of signalling and cytokine release and leads to platelet aggregation, as thrombocytopenia observed in this and previous^31^ studies in rats. Complement activation has been shown to induce production of platelet activating factor (PAF) and TxA_2_.^12^ Both of them are potent platelet activators that may induce platelet aggregation and microvascular occlusion, which is consistent with our finding of a partial occlusion of microvessels and reduced CSA. The results are in parallel with a previous study that PEG-MNPs, but not MNPs, may cause dynamic retention of agglomerate in microcirculation,^32^ and PEGylated nanoparticles may induce platelet aggregation.^33^ Since cyclooxygenase blockade by indomethacin exhibited no effect on the hemodynamics, our study does not support a role of TxA_2_ in the hemodynamic effects induced by PEG-MNPs in rats. The results are consistent with a previous finding that cyclooxygenase blockade by aspirin did not inhibit nanoparticle-induced platelet aggregation.^34^ Nevertheless, Yamanaka S. et al. has demonstrated that PAF (*i.v.*) induced similar hemodynamic responses in dogs, with hypotension, profound decrease in CO, and increase in vascular resistance;^35^ it was proposed that reduction in venous return might mediate the reduction in CO and arterial pressure, as observed in our study. Therefore, immobilized, but not free PEG may activate the complement system on the surface of the nanocomposites that triggers transient occlusion of microvasculature, as demonstrated by an increase in PVR. Hence, blood emerged into abdominal aorta, resulting in transient increase of ABF and ACSA before declining (Figure 1). Therefore, decreased venous return may play a key role in the hemodynamic responses. Nevertheless, whether PAF plays a critical role in PEG-MNP-induced hemodynamic changes requires further study.

Although the protein corona composition may be determined by a variety of parameters, including time of exposure, temperature, and static *vs.* dynamic conditions etc.,^36^ our results with the hard corona profiles are consistent with previous finding that the composition of the corona proteins may determine the characteristics of the nano-biointerface,^37^ and support the idea that bio-nano interaction may cause hemodynamic consequence.^31^ Although complement C3 has been found as a major component in the corona of a variety of nanoparticles with different coatings,^38^ our results suggest that PEG as part of the coating may greatly enhance complement C recruitment and probably activation on the surface of nanoparticles.

In our study, MNPs without PEGylation may prevent hemodynamic responses induced by PEG-MNPs. Proteomic study suggests that abundant plasma proteins were removed after the first exposure to both MNPs and PEG-MNPs (Figure 4). Therefore, depletion of complement proteins is likely contributed to tachyphylaxis upon second exposure. Our result is consistent with previous finding that CARPA-induced by PEGylated liposomal preparation Doxil® was prevented by pre-treatment of a placebo of PEGylated liposomes with the potential to induce complement activation.^39^

Dextran-coated iron oxide nanoparticles has been demonstrated safety without induction of acute HSRs.^40^ Although the surface characteristics may play a major role in complement activation and subsequent immune cell responses,^41^ PEG moiety is not the sole factor in the nano-bio interface. It has been shown that PEG on the polyurethane nanocomposites did not induce significant complement activation or pro-inflammatory cytokine release.^42^ Anti-PEG antibody may exist in the plasma of humans, but much less, if any, in naïve rats; complement activation is expected to be more vigorous in the presence of anti-PEG antibody,^43^ especially in the post-COVID-19 era.

As to endothelial function, our results indicate that two consecutive doses of MNPs or PEG-MNPs did not alter acute vasodilator response to Ach, suggesting these particles at such dose did not jeopardize endothelium-dependent vascular function *in vivo*. Although nanoparticle uptake by cultured endothelial cells has been well recognized and studied;^22, 23^ to our knowledge, no direct evidence indicate that endothelial uptake may occur in a healthy vessel under the flow/shear condition *in vivo*. Nevertheless, high dose of iron oxide nanoparticles (20 mg/kg, *i.v.*) may induce increase in eNOS activity measured 3 days after injection.^44^ The rat model appears to be able to respond to a direct vasodilator that increases blood flow; therefore, PEG-MNP-induced hypotension is unlikely mediated by a vasodilator effect.

## CONCLUSION

Hypotension induced by PEG-MNP in rats is associated with complement activation, tachyphylaxis, and thrombocytopenia, comparable to acute hypersensitivity induced by PEGylated nanomedicine in humans. This rat model demonstrated an HSR-associated increase in vascular resistance and a decrease in cardiac output; therefore, PEGylated MNP-induced-hypotension is likely mediated by a reduced venous return due to transitory occlusion of the microvessels. Both *in vitro* and *in vivo* results suggest that consumption or depletion of complement proteins may be contributed to tachyphylaxis induced by PEGylated nanocomposites. Our study reveals the detailed hemodynamic mechanism of HSR that may provide novel interpretations of the HSR-related symptoms, and offer new opportunities for developing novel strategies for preventing and treating HSR induced by PEGylated nanomedicine.

## NOVELTY AND SIGNIFICANCE

What is known?

➢ PEGylated nanocomposites may induce acute hypersensitivity reactions, including complement activation, hypotension and thrombocytopenia.
➢ PEGylated liposome-induced acute hypersensitivity reactions were associated with release of a variety of vasoactive cytokines, including histamine, platelet activating factor and thromboxane A_2_.

What new information does this article contribute?

➢ PEG-MNP-induced hypotension is associated with reduction in renal/cremaster flow and cardiac output, and increase in systemic vascular resistance in rats.
➢ The hard corona of PEG-MNP is composed of more complement proteins compared to that of pristine MNP, including C3, C6 and C9.
➢ Depletion of complements with cobra venom factor prevents PEG-MNP-induced hemodynamic effects in rats, suggesting a critical role of complement activation.

## ACKNOWLEDGMENT

The authors appreciate a gift of CVF from S. R. Roffler, Institute of Biomedical Sciences, Academia Sinica, Taiwan. The authors thank Proteomics Core Lab, Molecular Medicine Research Center, Chang Gung University for the proteomic analysis and Sonya Y. Hsueh for professional assistance with the graphic abstract.

## ASSOCIATED CONTENT

Supporting Information.

A PDF file supporting tables and figures is provided.

## AUTHOR INFORMATION

### Authorship Contributions

Conceptualization, Y.H.M., J.S.Y.; Methodology, M.T.S, K.Y.C.; Investigation, S.T.N., Y.C., S. Y.C., H.M.C.; Writing, S.T.N, Y.C., Y.H.M.; Supervision and fund acquisition, Y.H.M. The manuscript was written through the contributions of all authors. All authors have approved the final version of the manuscript. ‡These authors contributed equally.

## ABBREVIATIONS

ABF: aortic blood flow
AP: arterial pressure
AVR: aortic vascular resistance
CARPA: complement activation-related pseudoallergy
CO: cardiac output
CSA: cross-sectional area
CVF: cobra venom factor
HSR: hypersensitivity reactions
MAP: mean arterial pressure
MFP: microparticle-free plasma
MNP: magnetic nanoparticle
PEG: polyethylene glycol
PAF: platelet-activating factor
pI: isoelectric point
PVR: peripheral vascular resistance
RBF: renal blood flow
RVR: renal vascular resistance
SV: stroke volume
TxA_2_: thromboxane A_2_

